# Comparison of the microbiome and mycobiome in tissues of the tropical carnivorous epiphytic herb *Utricularia jamesoniana*

**DOI:** 10.1101/2023.09.09.556968

**Authors:** Valeria Naranjo-Aguilar, Rebeca Mora-Castro, Jessica Morera-Huertas, Rafael H. Acuña-Castillo, Keilor Rojas-Jimenez

## Abstract

*Utricularia jamesoniana*, a small epiphytic plant found in wet tropical forests, stands out for its carnivorous habit, intricate trap system, and small but beautiful and complex flowers. This species remains relatively understudied despite its wide geographical distribution and curious adaptations. In this study, we employed 16S rRNA and ITS sequencing to compare the prokaryotic and fungal communities within leaves and traps of *U. jamesoniana*. The analysis of amplicon sequence variants (ASVs) unveiled notable differences in community composition depending on the plant tissue and type of microorganism. Prokaryotic communities predominantly comprised Proteobacteria and Actinobacteriota, featuring genera such as *Acidocella, Bradyrhizobium, Ferritrophicum*, and *Ferrovum*. Fungal communities were dominated by Ascomycota and Basidiomycota, encompassing representatives of Dothideomycetes, Sordariomycetes, Eurotiomycetes, and Agaricomycetes, as well as ASVs related to Mycosphaerellaceae, *Colletotrichum, Aspergillus*, and *Thanatephorus*. We determined that the prokaryotic diversity has higher in the bladders with respect to the leaves. Fungal communities, in turn, were more diverse in leaves than in bladders. This study sheds light on the microbial communities associated with this carnivorous epiphyte and provides valuable insights into the intricate relationships between the plant and its microbial inhabitants across different tissues.

## 1 Introduction

Lentibulariaceae (Lamiales) is a family of carnivorous herbs composed of three genera: *Genlisea, Pinguicula*, and *Utricularia*. Among these, *Utricularia* is the most diverse, with terrestrial, epiphytic, lithophytic, rheophytic, and aquatic species. These plants are commonly referred to as “bladderworts” due to their complex traps, which are shaped like bladders. With approximately 250 species, *Utricularia* and similar species-rich *Drosera* (Droseraceae) are recognized as the most diverse genera of carnivorous plants (Fleischmann, 2015; Rutishauser, 2016). *Utricularia* is also the most widely distributed carnivorous plant genus, with a sub-cosmopolitan distribution encompassing every continent except Antarctica. The centers of diversity for *Utricularia* are Australia and the Neotropics (Taylor, 1989; Henning et al., 2021).

*Utricularia* has become a focus of increasing attention from the scientific community because the homologies of the organs that constitute their structure, compared to more typical plants, remain unclear (Rutishauser, 2016). Nonetheless, their vegetative structures are highly diverse and perform functions of rhizoids, rhizomes, stolons or pseudobulbs, and photosynthetic organs that resemble leaves (for practicality purposes, we will call these structures simply leaves, even if their homology has been subject to discussion). The stolons typically emerge from the base of the inflorescences and are robust in aquatic species but delicate and hidden in the substrate in terrestrial and epiphytic species. Furthermore, the genus includes some angiosperms with the smallest known genomes (Ibarra-Laclette et al., 2013).

Although most species of *Utricularia* are aquatic or inhabit wet soil, some members have evolved as epiphytes, particularly in tropical wet forests. Among these, several species of the Neotropical section Orchidioides, which includes 17 accepted species, are known to have such a habit (Henning et al., 2021). *Utricularia jamesoniana* (**Figure 1**) is one of the most widely distributed species in this monophyletic section and can be found from Mexico to Bolivia and northern Brazil, including Hispaniola and the Lesser Antilles (Gomes Rodrigues et al., 2017; Silva et al., 2018). It generally grows on tree trunks and branches, associated with bryophytes in humid to pluvial, frequently cloudy forests, including forest edges, secondary growth, and pastures (Crow, 2007; Valdés, 2008).

**Figure 1.**
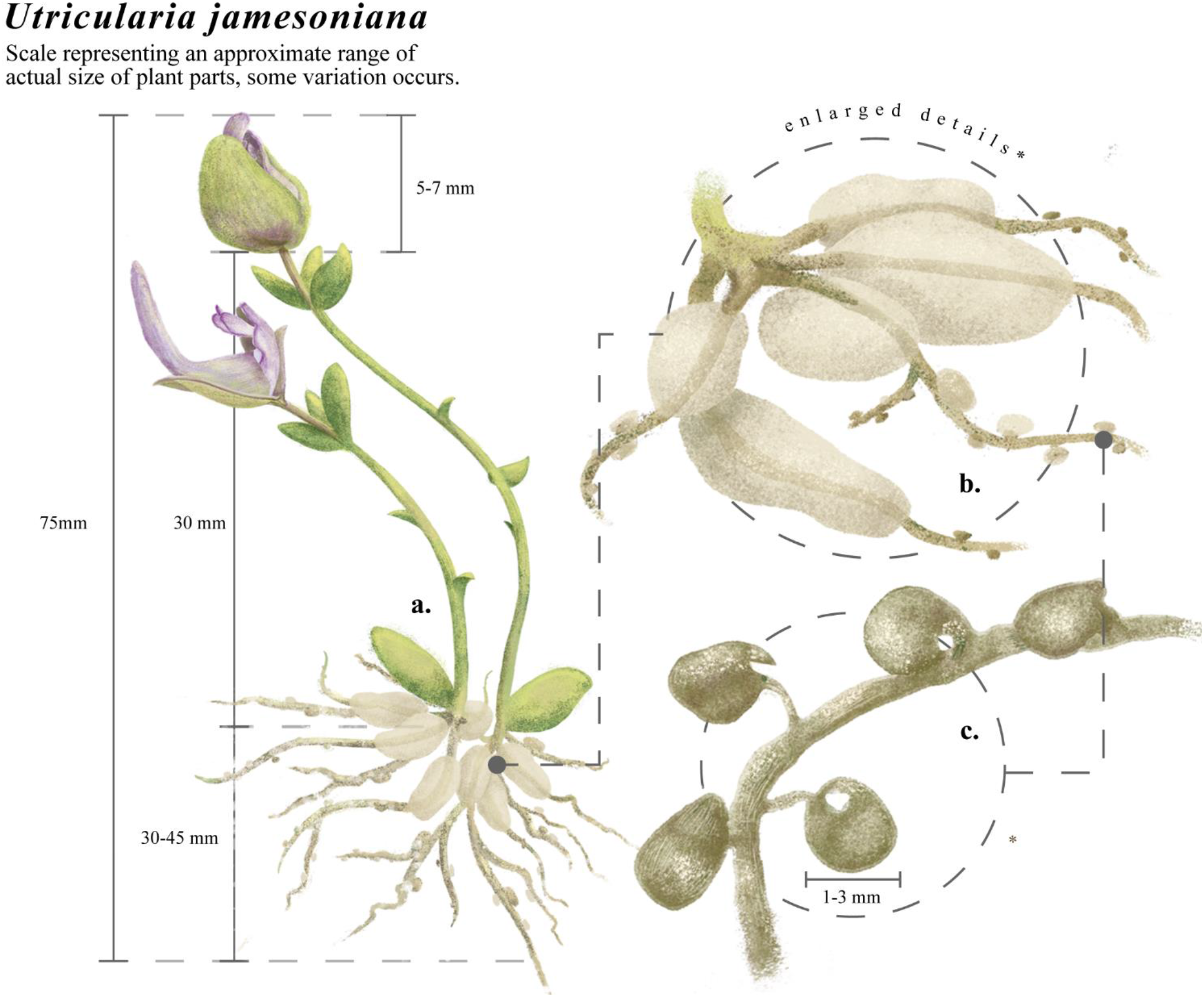
Utricularia jamesoniana. **A**. habit, **B**. stolons with tubers, **C**. traps. Measurements refer to the approximate sizes of different structures. Drawing prepared by Victor Adrian Mora Mora.

The bladders (or traps) have been reported in all *Utricularia* species (Miranda et al., 2021). They are seemingly foliar structures whose function is small prey capture through the secretion of enzymes and are considered to be the fastest carnivorous trap system in plants, although they have mainly been studied in aquatic species (Adamec et al., 2010; Westermeier et al., 2017). A mature trap is triggered by elastically loading the walls of the bladder as water is removed from the trap lumen, causing the walls to bend inward and generate a lower-than-ambient internal pressure. When prey touches the trigger hairs at the opening, the movable gate or valve bends inward, and the prey is drawn within a few milliseconds due to lower pressure within the bladder (Peroutka et al., 2008; Ortega Ardila and Romero Salgado, 2016).

In general, solid objects such as prey experience three forces that propel the prey toward the trap: pressure, viscous drag, and acceleration reaction. The prey is subsequently dissolved and digested by specialized cells. Traps can be ready to capture new prey again in 10-15 minutes, but water evacuation continues longer. It is worth noting, however, that the bladders of epiphytic species can differ significantly from those found in aquatic species, which may result in differences in their trapping mechanism (Givnish et al., 2018; Deban et al., 2020; Singh et al., 2020).

Numerous investigations have been conducted on the prey spectra of aquatic *Utricularia* (Walker, 2004; Sanabria-Aranda et al., 2006). However, less attention has been paid to exploring prey-capturing abilities in terrestrial species, in which their ability to catch prey has also been confirmed (Moseley, 1884; Jobson and Morris, 2001). The reported diet types include micro-crustaceans, nematodes, rotifers, insect larvae, tadpoles, and fish larvae (Sirová et al., 2003; Płachno et al., 2006; Martens and Grabow, 2011).

Several different clades of microorganisms that inhabit the traps have been documented, including bacteria, ciliates, algae, and other protozoans. These microorganisms play crucial roles in the conversion of decaying matter. Certain species are considered commensals, while others are recognized as parasites, further adding to the complexity of the bladder ecosystem (Płachno et al., 2006; Peroutka et al., 2008; Alkhalaf et al., 2009; Adamec, 2011; Ibarra-Laclette et al., 2013; Sirová et al., 2018b). Furthermore, the microbial communities in the bladders can vary depending on the anatomical structure of the trap as well as the stage of development of the plant (Friday, 1989; Płachno et al., 2012; Sirová et al., 2018a).

Information on this species of epiphytic carnivorous plant is limited, except for some taxonomic treatments providing basic descriptions of the morphology, ecology, and distribution (Oliver, 1860; Taylor, 1989; Crow, 2007; Guedes et al., 2023). To our knowledge, no studies have explored the prokaryotic microbiota and mycobiome in tissues of *Utricularia jamesoniana*, including its traps. In this work, we provide the first characterization of the bacterial and fungal communities associated with the bladders and leaves of *Utricularia jamesoniana*, which contribute to a better understanding of the physiology and ecology of this carnivorous species.

## 2 Materials and Methods

### 2.1 Sample collection

In July 2022, we collected representative *Utricularia jamesoniana* plants (**Figure 1**) from the Robert and Catherine Wilson Botanical Garden, part of Las Cruces Biological Station (LCBS) owned by the Organization for Tropical Studies (OTS). LCBS is in southern Costa Rica in the Coto Brus county of Puntarenas Province, ca. 5 km away in a straight line from the Panamanian border and 4 km in a straight line to the SW of the town of San Vito. The Garden is at about 1200 m asl on the Fila Cruces, part of the Fila Costeña Sur, located south of the Coto Brus Valley. The Garden is in the Premontane Rain Forest Life zone, according to Tosi (1969), and is surrounded by young and mature secondary forests. *Utricularia jamesoniana* is common in the garden grounds and can grow as an epiphyte (or facultatively epilithic) among dense bryophyte mats. The collected plants grew on the mossy stilt roots of mature, cultivated *Socratea exorrhiza* palms. Plant samples were placed in polyethylene bags, while a voucher was adequately preserved a 70% ethanol solution in the spirit collection of the USJ herbarium (*Acuña et al. 3183*). Plants were collected, transported to the laboratory, and processed on the same day. The plant collection was carried out with the respective permits of the authorities (collection License ACLAP-077-2021) and the official permission of the Biodiversity Commission of the Universidad de Costa Rica (CBio-73-2022, resolution #371).

### 2.2 Molecular analyses

Once in the laboratory, plant samples were washed thoroughly in running water, and healthy and physically undamaged tissues (leaves and bladders) were selected and sterilized on the surface with 70% ethanol and aseptically cut. The DNA was extracted from approximately 100 mg of the plant tissues using a DNA isolation kit (PowerSoil, Qiagen, USA) following the manufacturer’s instructions. The prokaryotic amplicon library was constructed based on the V4 hypervariable region of the 16S rRNA gene using universal primers 515F and 806R (Caporaso et al., 2011). The eukaryotic amplicon library was generated by amplifying the ITS2 region using primers ITS3-2024F (5’ GCATCGATGAAGAACGCAGC) and ITS4-2409R (5’ TCCTCCGCTTATTGATATGC). PCR amplification of targeted regions was performed using specific primers connecting with barcodes. The PCR products with the proper size were selected by 1% agarose gel electrophoresis. Each sample’s exact amount of PCR products was pooled, end-repaired, A-tailed, and further ligated with Illumina adapters. Libraries were sequenced on a paired-end Illumina platform to generate 250bp paired-end raw reads (Illumina Novaseq, Novogene Bioinformatics Technology Co., Ltd, CA, USA).

### 2.3 Biofinformatic analyses

We used the DADA2 version 1.21 to process the Illumina-sequenced paired-end fastq files and to generate a table of amplicon sequence variants (ASVs), which are higher-resolution analogs of the traditional OTUs (Callahan et al., 2016). Briefly, we removed primers and adapters, inspected the quality profiles of the reads, filtered and trimmed sequences with a quality score < 30, estimated error rates, modeled and corrected amplicon errors, and inferred the sequence variants. Then, we merged the forward and reverse reads to obtain the full denoised sequences, removed chimeras, and constructed the ASV table. We assigned taxonomy to the ASVs with the function *assignTaxonomy*, of DADA2. For the prokaryotic assignment, we used as input the SILVA reference database version 138.1 (Quast et al., 2013). We carried out a second taxonomic assignment of the prokaryotic ASVs using the tool IDTAXA of DECIPHER (Murali et al., 2018) with the same version of the SILVA database and using the RDP database version 18 (http://rdp.cme.msu.edu/). For the fungal taxonomic assignment, we used the UNITE ITS database version 8.3 (https://doi.org/10.15156/BIO/1280049). We carried out a second taxonomic assignment of the fungal ASVs using the tool IDTAXA of DECIPHER with the same version of UNITE as a reference and a confidence threshold >60%, and additionally, a third classification using the Classifier tool (Wang et al., 2007) implemented in the Ribosomal Database Project (http://rdp.cme.msu.edu/) using as reference the Warcup Fungal ITS trainset V2 database (Deshpande et al., 2015). We verified and manually curated the consistency between the taxonomic assignments of the different programs for prokaryotes and fungi. In cases of discrepancies, a comparison with the BLAST tool of NCBI Genbank was applied. Sequence data were deposited at the NCBI Sequence Read Archive under accession number PRJNA1005337 (https://www.ncbi.nlm.nih.gov/sra/PRJNA1005337). In the 16S rRNA prokaryotic dataset, we removed sequences assigned to Chloroplast and Eukarya. This process generated 518.491 sequences from the four samples. The average number of sequences per sample was 129.623 (ranging from 111.131 to 149.377). In the ITS2 eukaryotic dataset, we removed sequences assigned to vertebrates and plants. This process generated 283.135 sequences from the samples. The average number of sequences per sample was 70.784 (ranging from 57,195 to 94.867).

### 2.4 Statistical analyses

The statistical analyses and visualization of results were performed with the R statistical program (R Core Team, 2023) and the Rstudio interface. Package Vegan v2.6-4 (Oksanen et al., 2022) was used to calculate alpha diversity estimators and non-metric multidimensional scaling analyses (NMDS). Data tables with the amplicon sequence variants (ASV) abundances were normalized into relative abundances and converted into a Bray–Curtis similarity matrix. To determine significant differences between the bacterial and fungal community composition according to the plant tissue, we used the non-parametric multivariate analysis of variance (PERMANOVA) and pairwise PERMANOVA (adonis2 function with 999 permutations). To estimate differences in the relative proportion of microorganisms and the diversity indices, we used the non-parametric Kruskal-Wallis test, while pairwise comparisons were performed using Wilcox and Dunn’s tests (with Bonferroni adjustment).

## 3. Results

In this work, we compare the prokaryotic and fungal communities in the leaves and bladders of the carnivorous epiphytic plant *Utricularia jamesoniana*. The prokaryotic community of the samples was composed of 2.258 amplicon sequence variants (ASVs), according to the analysis of sequences of the V4 region of the 16S rRNA gene. All the prokaryotic sequences were assigned to 44 phyla and 87 classes. Proteobacteria were the most abundant, representing 66% of all sequences and 37.5% of the ASVs. Actinobacteriota represented 11.3 % of the sequences and 9.3% of the ASVs, and Acidobacteriota represented 3.9% of the sequences and 5.6% of the ASVs. Other abundant groups were Firmicutes, Verrucomicrobiota, Planctomycetota, and Cyanobacteria. Archaea represented only 0.28% of the sequences and 0.56% of the ASVs.

The fungal community was composed of 1.620 amplicon sequence variants, according to the analysis of sequences of the ITS region. All the fungal sequences were assigned to 10 phyla and 32 classes. Ascomycota was the most abundant phylum comprising 79.6% of the sequences and 78.9% of the ASVs, while Basidiomycota represented 20.2 % of the sequences and 19.3% of the ASVs. Other less abundant fungal phyla included Rozellomycota and Chytridiomycota.

The fungal classes with the highest relative abundance were Dothideomycetes, Sordariomycetes, Eurotiomycetes, and Agaricomycetes, which aligns with the fungal community structure described for other carnivorous plants of the same family (Rueda-Almazán et al., 2021). Within Dothideomycetes, the most abundant ASVs were unclassified members of Mycosphaerellaceae and Cladosporiaceae, while *Colletotrichum, Aspergillus*, and *Thanatephorus* were the most abundant genera within Sordariomycetes, Eurotiomycetes, and Agaricomycetes, respectively.

The bacterial classes with the highest relative abundance were Alphaproteobacteria and Gammaproteobacteria. The most abundant genera within Alphaproteobacteria were *Acidocella, Bradyrhizobium*, and *Rhizobium*. Rhizobial bacteria have been reported as one of the most abundant groups in some aquatic *Utricularia* species, while *Acidocella* and *Bradyrhizobium* have also been reported in the traps of different carnivorous plants (Sickel et al., 2019; Grothjan and Young, 2022). The most abundant genera within Gammaproteobacteria were *Ferritrophicum, Ferrovum*, and *Chromobacterium*. From these, only *Chromobacterium* has been described as part of the microbiome of any *Utricularia* (Grothjan and Young, 2022). In general, we observed differences in the composition of some particular groups of the prokaryotic communities between the leaves and traps (**Figure 2A**).

**Figure 2.**
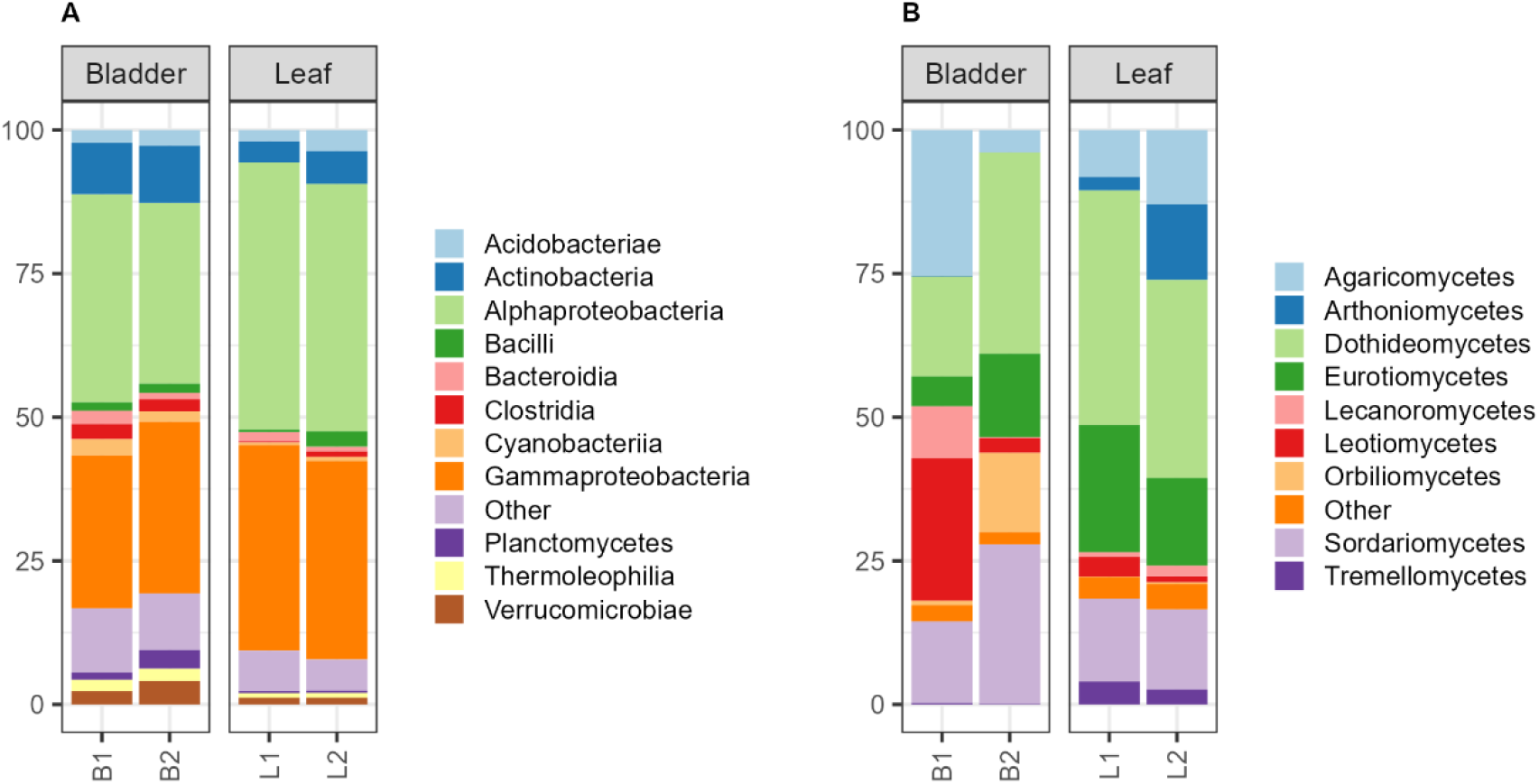
Relative abundance of the main taxonomic groups in bladders and leaves of *Utricularia jamesoniana*. Panel **A** corresponds to the main prokaryotic groups, and Panel **B** to the main fungal groups.

We also observed variations in the composition of the fungal communities within samples and between tissues. Notably, we detected a higher proportion of Lecanoromycetes, Leotiomycetes, Orbiliomycetes, and Sordariomycetes in the bladder, while Dothideomycetes and Eurotiomycetes were more abundant in the leaves. We detected 25 and 30 classes of fungi in the bladder and leaf tissues, respectively. Chytridiomycetes were detected only in the bladder, where they have been found actively growing in aquatic *Utricularia* species (Sirová et al., 2018b). Conversely, some classes were only present in the leaves, including Agaricostilbomycetes, Saccharomycetes, and Rhizophydiomycetes (**Figure 2B**).

We observed notable differences in the alpha diversity indices of prokaryotic communities in the tissues of *Utricularia jamesoniana*. The richness of prokaryotic ASVs has higher in the bladders with respect to the leaves, with mean richness values of 1174 and 356, respectively. On the other hand, the mean value of the Shannon diversity index in the traps was also higher than that of the leaves, with mean values of 6.0 and 4.3, respectively (**Figure 3**). We also observed differences in the indices of fungal communities. However, the mycobiome was more diverse in the leaves than in the traps. The richness of fungal ASVs has higher in the leaves with respect to the traps, with mean richness values of 652 and 433, respectively. On the other hand, the mean value of the Shannon diversity index in the leaves was also higher than that of the traps, with mean values of 4.2 and 3.7, respectively (**Figure 3**).

**Figure 3.**
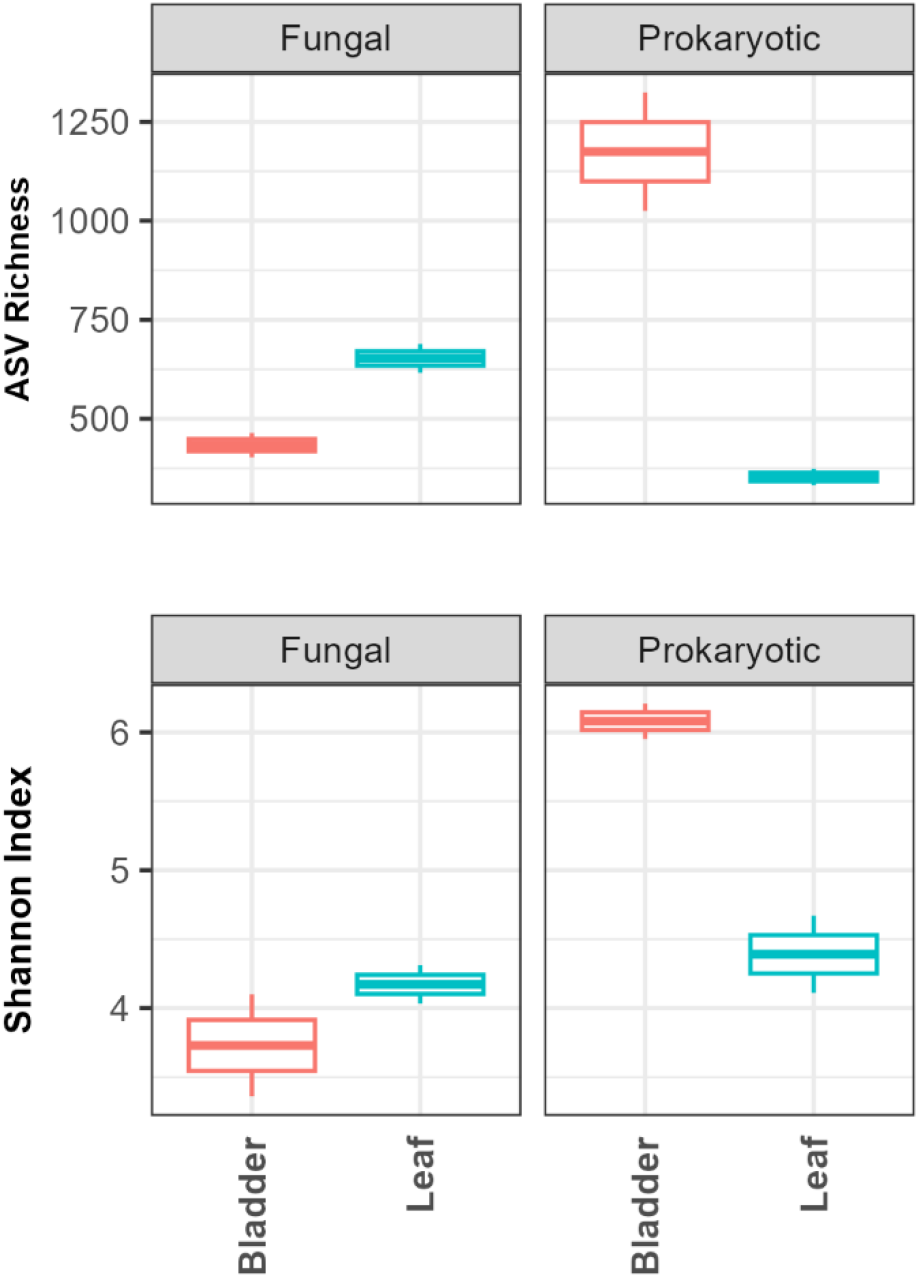
Alpha diversity values of prokaryotic and fungal communities in the bladder and leaves of *Utricularia jamesoniana*. The upper panel represents the richness of Amplicon Sequence Variants (ASVs), while the panel below shows the values of the Shannon Index.

The NMDS analysis of the bacterial communities showed that the trap samples clustered together and their community composition separated from the plant leaves (**Figure 4A**). However, the Permanova showed no significant differences in the structure of prokaryotic communities between tissues (Permanova > 0.05). The analysis of the fungal communities showed that the composition in both bladders and leaves tissues in one of the plants clustered together, whereas they separated in the second plant (**Figure 4B**). The permanova showed no significant differences in the structure of prokaryotic communities between tissues (Permanova > 0.05).

**Figure 4.**
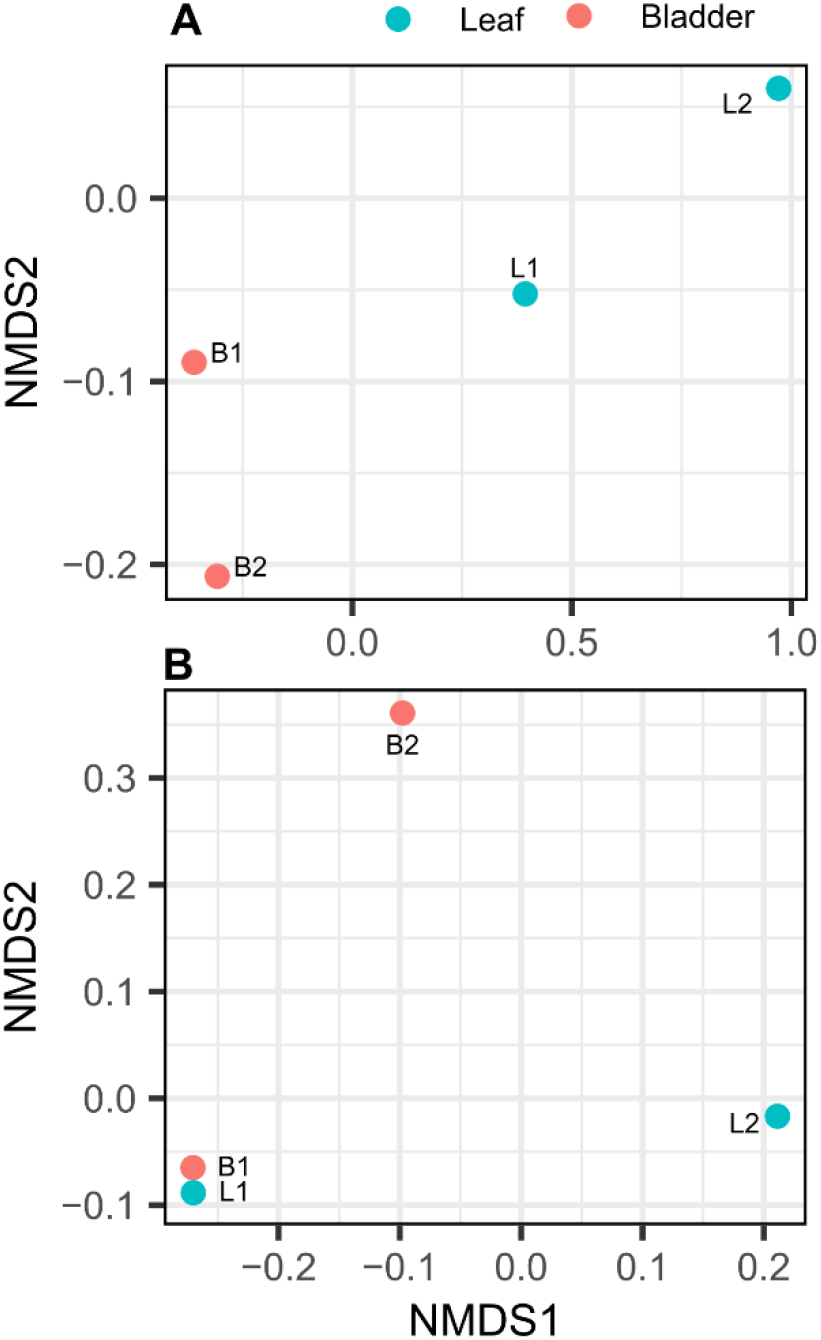
NDMS analysis of the microbial communities in bladders and leaves of *Utricularia jamesoniana*. Panel **A** corresponds to the prokaryotic groups, and Panel **B** to the fungal groups.

The heat map (**Figure 5A**) shows the variations in the relative abundances of the sample’s most abundant genera of prokaryotes. In general, most genera presented relative abundances of less than 5%. However, some of the most abundant genera in the bladder were *Bradyrhizobium, Ferritrophicum, Acidocella, Methylocapsa*, and *Mycobacterium*, while in the leaves, they were *Acidocella, Ferrovum, Methylocapsa, Rhizobium*, and *Gallionella. Acidocella* and *Methylocapsa*.

**Figure 5.**
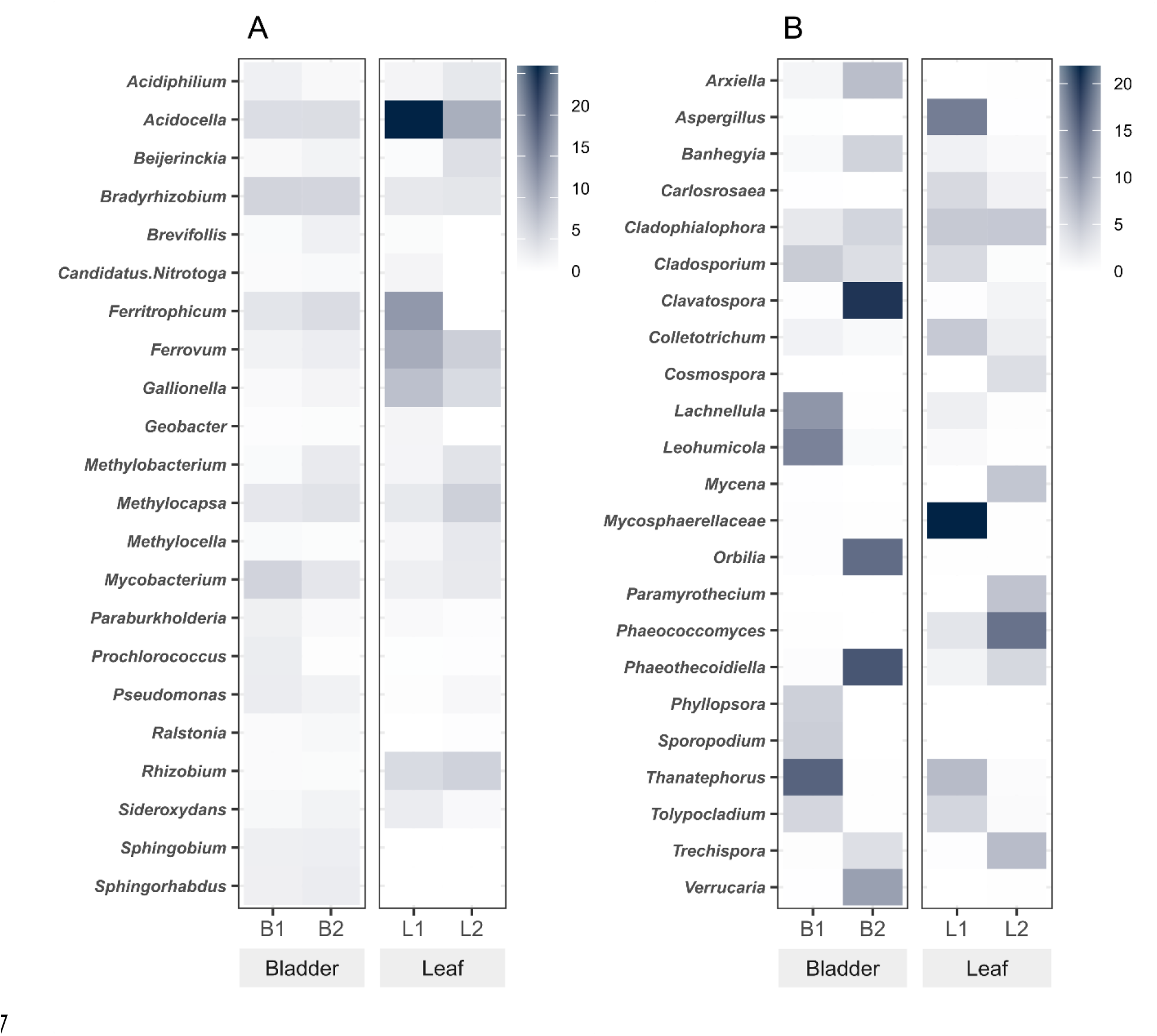
Heatmap of the most abundant genera in the bladder and leaves tissues of *Utricularia jamesoniana*. Panel **A** corresponds to the prokaryotic genera determined according to the relative abundance of 16S rRNA gene sequences, and Panel **B** to the fungal genera determined according to the relative abundance of ITS sequences.

Our results confirm the previously observed differences in fungal diversity within and between samples regarding the most prevalent fungal genera across the tissue samples. In particular, we identified several fungal genera that were highly abundant in traps, including *Thanatephorus, Clavatospora*, and *Phaeothecoidiella*. In contrast, leaves’ most prevalent fungal genera were an unclassified member of the Mycosphaerellaceae, *Aspergillus*, and *Phaeococcomyces*.

## 4 Discussion

This study elucidates the distinctive differentials in prokaryotic and fungal communities between the traps of *Utricularia jamesoniana* and its leaves. The specialized bladders or traps constitute a singular ecological niche that harbors a multitude of microbial communities. These communities potentially exert a significant influence on plant fitness by participating in prey digestion and facilitating environmental adaptation (Caravieri et al., 2014; Sirová et al., 2018a). Notably, the resident microbiome within these traps may undergo modifications due to the introduction of microbial communities from captured prey, as well as the assimilation of metabolizable compounds derived from the prey. It is pertinent to recognize that each prey item might harbor its own distinct microbial diversity, as well as inherent chemical characteristics, thereby introducing an additional layer of complexity to the trap’s microbiome composition.

The predominance of *Bradyrhizobium*, a nitrogen-fixing bacterium (Zahran, 1999), was particularly conspicuous within the traps. Nevertheless, uncertainty persists regarding the nitrogen-fixing capability of this species within a non-root nodule, such as the bladder. On the other hand, it is plausible that *Bradyrhizobium* might serve additional functions, such as metabolizing different carbon sources (VanInsberghe et al., 2015). Furthermore, the traps exhibited a noteworthy abundance of metal-oxidizing bacteria such as *Acidocella, Ferritrophicum*, and *Ferrovum* which suggests the occurrence of these processes within this plant tissue (Weiss et al., 2007; Johnson et al., 2014; Okamoto et al., 2017). This assemblage of microbial species suggests an environment conducive to stimulating microbial metabolism, leading to the release and recycling of nutrients (Sirová et al., 2003, 2018a).

The interior of the trap is completely sealed and maintains an anoxic environment (Adamec, 2007), and hosting a complex microbial food web responsible for a significant proportion of enzymatic activity associated with prey digestion (Sirová et al., 2009; Caravieri et al., 2014). The preservation of anoxic conditions within the traps potentially serves as the mechanism for incapacitating captured prey (Adamec, 2007).

Given this context, it stands to reason that the microbial community dwelling within these traps has adapted to this unique environment, which experiences intermittent spikes of elevated oxygen concentration subsequent to firing events, followed by periods of anoxia (Adamec, 2007). These fluctuating conditions create a niche favoring the coexistence of organisms that exhibit heightened tolerance to anoxia (Richards, 2001; Sirová et al., 2003), consequently contributing to prey degradation (Caravieri et al., 2014). These conditions could also be responsible for the changes in the composition of the fungal communities in this tissue and the lower values of diversity observed.

Proteobacteria emerged as the predominant inhabitants within the *Utricularia jamesoniana* tissues. The prevalence and roles of this taxonomic group are intricately linked to the microenvironment provided by the plant tissue, its genetic composition, physiological attributes, as well as the biotic and abiotic factors inherent in the surrounding environment (Hardoim et al., 2015; Brader et al., 2017; Afzal et al., 2019). Prior microbiome investigations associated with other carnivorous plants have consistently pinpointed Proteobacteria as the most abundant phylum (Alcaraz et al., 2016; Chan, 2019; Sickel et al., 2019).

Taxonomic parallels with earlier studies conducted on other aquatic *Utricularia* indicate the prevalence of *Rhizobium* as one of the most abundant genera within certain species. Likewise, *Acidocella* and *Bradyrhizobium* have been noted in the traps of diverse carnivorous plants (Sickel et al., 2019; Grothjan and Young, 2022). Among Gammaproteobacteria, *Chromobacterium* has also been described as part of the microbiome across several *Utricularia* species (Grothjan and Young, 2022).

One of the most revealing results of this study was the elevated levels of prokaryotic species richness and Shannon diversity detected within the traps, constituting a digestive ecosystem (Adamec, 2007; Sirová et al., 2018b, 2018a) where microorganisms face challenging living conditions typical of a digestive ecosystem. This intriguing observation prompts speculation that the increased richness and diversity of communities within the traps stem from the dynamic nature of this environment. In contrast, the conditions within the leaves can be more stable.

The diversity of the endophytic fungal community within the traps exhibited lower values compared to that observed in the leaves. This trend stands in contrast to the patterns observed within bacterial communities. Notably prevalent within this tissue were the genera *Thanatephorus, Leohumicola, Clavatospora, Orbilia*, and *Phaeothecoidiella*. However, the distinctive attributes and mechanisms underlying these genera’s apparent compatibility within the host traps fluctuating anaerobic microenvironment demand further exploration. Particularly intriguing is the prevalence of *Thanatephorus*, known as the teleomorph of *Rhizoctonia*, a recognized plant pathogen (Donk, 1956; Anderson, 1982), which was present even when no disease symptoms were observed on the plants examined.

In general, the phylum Ascomycota predominated in the tissues of *Utricularia jamesoniana*, a pattern consistent with earlier findings in other carnivorous plants belonging to the same botanical family (Rueda-Almazán et al., 2021). Particularly noteworthy is that a substantial proportion, approximately 45%, of the fungal genera were present in both the traps and leaves. This observation underscores the notion that certain endophytic fungi can colonize diverse tissues within the same plant organism effectively (Wu et al., 2013; Behie et al., 2015; Rueda-Almazán et al., 2021).

The interactions between plants and endophytes have been recognized for their myriad benefits to plants, particularly species that thrive in challenging or nutrient-deficient environments, as is the case with carnivorous plants (Ellison and Adamec, 2018; Jobson et al., 2018; Rueda-Almazán et al., 2021). However, It is crucial to highlight that not all identified fungi necessarily qualify as pure endophytes, as some could originate from external and potentially serve as nutrient sources (Sirová et al., 2018b).

In the present investigation, we identified distinct groups of genuine endophytes, such as *Aspergillus, Cladosporium, Colletotrichum*, and Mycosphaerellacea. These groups have previously been isolated from both leaves and traps in other carnivorous plant species (Quilliam and Jones, 2010; Glenn and Bodri, 2012; Lee et al., 2014; Naseem and Kayang, 2018; Rueda-Almazán et al., 2021). However, some genera were exclusive to either the traps or leaves. *Pseudochaetosphaeronema, Periconia, Remotididymella*, and *Phragmotaenium* have also been described for the first time in an epiphytic carnivorous plant species. These findings underscore the distinctive biodiversity ecosystem hosted by *Utricularia jamesoniana*, justifying comprehensive further investigation.

Finally, the exploration of plant-microbial interactions serves as a window into the intricate web of ecological relationships spanning diverse lineages and their evolutionary trajectories. The dynamic composition of these microbial communities can fluctuate in response to a spectrum of biotic and abiotic factors that shape these interactions, where microorganisms undertake an array of functions, encompassing enzymatic activities such as prey digestion, fortification against pathogens, facilitation of growth, and protection (Lee et al., 2014; Hung and Lee Rutgers, 2016; Rueda-Almazán et al., 2021). Examining the microbiome and mycobiome of carnivorous plants provides a captivating model that advances our understanding of plant ecology, evolution, and the intricate interplay between plants and their microbial inhabitants.

We contribute novel insights that enhance the comprehension of the microbiological community associated with *Utricularia jamesoniana*, shedding light on previously undocumented groups present within plant leaves and traps. For future investigations, it is imperative to incorporate a larger sample size to facilitate robust statistical analyses while exercising caution to prevent any adverse impact on the vulnerable populations of this delicate species. Furthermore, it is necessary to explore in more detail the physiological and biochemical nuances characterizing the traps. This entails discerning the intrinsic microbiome residing within the traps and disentangling it from the influx of nutrient-rich biological material. Additionally, there is a need to meticulously document any discernible shifts in microbial populations triggered by the plant’s climatic seasonality and phenological state.

## 5 Conclusions

This study offers insight into the intricate microbial and fungal communities associated with traps and leaves of *Utricularia jamesoniana*. The distinctive biodiversity composition suggests plausible ecological connections between the plant tissues and the microbiome. These interactions and their potential entanglement with the fungal communities hold promise for illuminating the intricate host-endophyte dynamics. A more exhaustive microbiome characterization can unveil the plant’s dietary preferences, nutrient cycling potential, and the spectrum of associated microorganisms. The description of several previously unreported fungal groups highlights the specific and under-explored nature of this microbial and fungal habitat. We emphasize the need for further investigation to unravel the microbiome and mycobiome of this epiphytic carnivorous plant, its intricate relations, and interaction dynamics.

## 6 Conflict of Interest

The authors declare that the research was conducted in the absence of any commercial or financial relationships that could be construed as a potential conflict of interest.

## 7 Author Contributions

VNA, RMC and KRJ contributed to the conception and design of the study. RAC, JMH and VNA contributed to investigation and data analysis. KRJ wrote the first draft of the manuscript. RMC supervised the project. All authors contributed to the manuscript’s revision and read, and approved the submitted version.

## 8 Funding

This project was partially supported by Vicerrectoría de Investigación, University of Costa Rica (project C2-234).

## 9 Acknowledgments

We thank Víctor Adrián Mora Mora for the design and drawing of Figure 1.

## 10 Data Availability Statement

The datasets presented in this study can be found in the NCBI Sequence Read Archive under accession PRJNA1005337: https://www.ncbi.nlm.nih.gov/sra/PRJNA1005337.

## Notes

### Competing Interest Statement

The authors have declared no competing interest.

## References

Adamec, L. (2007). Oxygen Concentrations Inside the Traps of the Carnivorous Plants Utricularia and Genlisea (Lentibulariaceae). Ann. Bot. 100, 849–856. doi:10.1093/aob/mcm182.

Adamec, L. (2011). Functional characteristics of traps of aquatic carnivorous Utricularia species. Aquat. Bot. 95, 226–233. doi:10.1016/j.aquabot.2011.07.001.

Adamec, L., Sirová, D., Vrba, J., and Rejmánková, E. (2010). Enzyme production in the traps of aquatic Utricularia species. Biologia (Bratisl). 65, 273–278. doi:10.2478/s11756-010-0002-1.

Afzal, I., Shinwari, Z. K., Sikandar, S., and Shahzad, S. (2019). Plant beneficial endophytic bacteria: Mechanisms, diversity, host range and genetic determinants. Microbiol. Res. 221, 36–49. doi:10.1016/j.micres.2019.02.001.

Alcaraz, L. D., Martínez-Sánchez, S., Torres, I., Ibarra-Laclette, E., and Herrera-Estrella, L. (2016). The Metagenome of Utricularia gibba’s Traps: Into the Microbial Input to a Carnivorous Plant. PLoS One 11, e0148979. doi:10.1371/JOURNAL.PONE.0148979.

Alkhalaf, I. A., Hübener, T., and Porembski, S. (2009). Prey spectra of aquatic Utricularia species (Lentibulariaceae) in northeastern Germany: The role of planktonic algae. Flora -Morphol. Distrib. Funct. Ecol. Plants 204, 700–708. doi:10.1016/j.flora.2008.09.008.

Anderson, N. A. (1982). The genetics and pathology of Rhizoctonia solani. Annu. Rev. Phytopathol. 20, 329–347.

Behie, S. W., Jones, S. J., and Bidochka, M. J. (2015). Plant tissue localization of the endophytic insect pathogenic fungi Metarhizium and Beauveria. Fungal Ecol. 13, 112–119. doi:10.1016/J.FUNECO.2014.08.001.

Brader, G., Compant, S., Vescio, K., Mitter, B., Trognitz, F., Ma, L.-J., et al. (2017). Ecology and Genomic Insights into Plant-Pathogenic and Plant-Nonpathogenic Endophytes. Annu. Rev. Phytopathol. 55, 61–83. doi:10.1146/annurev-phyto-080516-035641.

Callahan, B. J., McMurdie, P. J., Rosen, M. J., Han, A. W., Johnson, A. J. A., and Holmes, S. P. (2016). DADA2: High-resolution sample inference from Illumina amplicon data. Nat. Methods 13, 581–583. doi:10.1038/nmeth.3869.

Caporaso, J. G., Lauber, C. L., Walters, W. A., Berg-Lyons, D., Lozupone, C. A., Turnbaugh, P. J., et al. (2011). Global patterns of 16S rRNA diversity at a depth of millions of sequences per sample. Proc. Natl. Acad. Sci. U. S. A. 108, 4516–4522. doi:10.1073/pnas.1000080107.

Caravieri, F. A., Ferreira, A. J., Ferreira, A., Clivati, D., de Miranda, V. F. O., and Araújo, W. L. (2014). Bacterial community associated with traps of the carnivorous plants Utricularia hydrocarpa and Genlisea filiformis. Aquat. Bot. 116, 8–12. doi:10.1016/J.AQUABOT.2013.12.008.

Chan, X. Y. (2019). Microbial diversity and bacterial biocatalytic activities in pitcher fluid of Nepenthes sp. Available at: http://studentsrepo.um.edu.my/id/eprint/12977.

Crow, G. E. (2007). “Lentibulariaceae,” in Manual de Plantas de Costa Rica, Volumen VI, Dicotiledóneas (Haloragaceae–Phytolaccaceae), eds. B. E. Hammel, M. H. Grayum, C. Herrera, and N. Zamora (Monographs in Systematic Botany from the Missouri Botanical Garden), 189–197.

Deban, S. M., Holzman, R., and Müller, U. K. (2020). Suction Feeding by Small Organisms: Performance Limits in Larval Vertebrates and Carnivorous Plants. Integr. Comp. Biol. 60, 852–863. doi:10.1093/icb/icaa105.

Deshpande, V., Wang, Q., Greenfield, P., Charleston, M., Porras-Alfaro, A., Kuske, C. R., et al. (2015). Fungal identification using a Bayesian classifier and the Warcup training set of internal transcribed spacer sequences. Mycologia, 14–293. doi:10.3852/14-293.

Donk, M. A. (1956). NOTES ON RESUPINATE HYMENOMYCETES-II* The tulasnelloid fungi. Reinwardtia 3, 363–379.

Ellison, A., and Adamec, L. (2018). Carnivorous plants: physiology, ecology, and evolution.

Fleischmann, A. (2015). Taxonomic Utricularia news. Carniv. Plant Newsl. 44, 13–16.

Friday, L. E. (1989). Rapid turnover of traps in Utricularia vulgaris L. Oecologia 80, 272–277. doi:10.1007/BF00380163.

Givnish, T. J., Sparks, K. W., Hunter, S. J., and Pavlovič, A. (2018). Why are plants carnivorous? Cost/benefit analysis, whole-plant growth, and the context-specific advantages of botanical carnivory. Oxford University Press doi:10.1093/oso/9780198779841.003.0018.

Glenn, A., and Bodri, M. S. (2012). Fungal Endophyte Diversity in Sarracenia. PLoS One 7, e32980. doi:10.1371/JOURNAL.PONE.0032980.

Gomes Rodrigues, F., Franco Marulanda, N., Silva, S. R., Płachno, B. J., Adamec, L., and Miranda, V. F. O. (2017). Phylogeny of the ‘orchid-like’ bladderworts (gen. Utricularia sect. Orchidioides and Iperua: Lentibulariaceae) with remarks on the stolon–tuber system. Ann. Bot. 120, 709–723. doi:10.1093/aob/mcx056.

Grothjan, J. J., and Young, E. B. (2022). Bacterial Recruitment to Carnivorous Pitcher Plant Communities: Identifying Sources Influencing Plant Microbiome Composition and Function. Front. Microbiol. 13, 791079. doi:10.3389/FMICB.2022.791079/BIBTEX.

Guedes, F. M., Gonella, P.M. Domínguez, Y., Moreira, A. D. R., Silva, S. R., Díaz, Y.C.A. Fleischmann, A. Menezes, C. G., Rivadavia, F., et al. (2023). Utricularia in Flora e Funga do Brasil. Jard. Botânico do Rio Janeiro. Available at: https://floradobrasil.jbrj.gov.br/FB19291.

Hardoim, P. R., van Overbeek, L. S., Berg, G., Pirttilä, A. M., Compant, S., Campisano, A., et al. (2015). The Hidden World within Plants: Ecological and Evolutionary Considerations for Defining Functioning of Microbial Endophytes. Microbiol. Mol. Biol. Rev. 79, 293–320. doi:10.1128/MMBR.00050-14.

Henning, T., Allen, J. P., and Rodríguez Rodríguez, E. F. (2021). A new species of Utricularia Sect. Orchidioides (Lentibulariaceae) from the Amotape-Huancabamba Zone of North Peru. Darwiniana, nueva Ser. 9, 299–311. doi:10.14522/darwiniana.2021.92.955.

Hung, R., and Lee Rutgers, S. (2016). “Applications of Aspergillus in Plant Growth Promotion,” in New and Future Developments in Microbial Biotechnology and Bioengineering (Elsevier), 223–227. doi:10.1016/B978-0-444-63505-1.00018-X.

Ibarra-Laclette, E., Lyons, E., Hernández-Guzmán, G., Pérez-Torres, C. A., Carretero-Paulet, L., Chang, T.-H., et al. (2013). Architecture and evolution of a minute plant genome. Nature 498, 94–98. doi:10.1038/nature12132.

Jobson, R. W., Baleeiro, P. C., and Guisande, C. (2018). Systematics and evolution of Lentibulariaceae: III. Utricularia. Oxford University Press doi:10.1093/oso/9780198779841.003.0008.

Jobson, R. W., and Morris, E. C. (2001). Feeding ecology of a carnivorous bladderwort (Utricularia uliginosa, Lentibulariaceae). Austral Ecol. 26, 680–691. doi:10.1046/j.1442-9993.2001.01149.x.

Johnson, D. B., Hallberg, K. B., and Hedrich, S. (2014). Uncovering a Microbial Enigma: Isolation and Characterization of the Streamer-Generating, Iron-Oxidizing, Acidophilic Bacterium “Ferrovum myxofaciens.” Appl. Environ. Microbiol. 80, 672–680. doi:10.1128/AEM.03230-13.

Lee, J., Tan, W., and Ting, A. (2014). Revealing the antimicrobial and enzymatic potentials of culturable fungal endophytes from tropical pitcher plants (Nepenthes spp.). Mycosphere 5, 364–377. doi:10.5943/mycosphere/5/2/10.

Martens, A., and Grabow, K. (2011). Early stadium damselfly larvae (Odonata: Coenagrionidae) as prey of an aquatic plant, Utricularia australis. Int. J. Odonatol. 14, 101–104. doi:10.1080/13887890.2011.568191.

Miranda, V. F. O., Silva, S. R., Reut, M. S., Dolsan, H., Stolarczyk, P., Rutishauser, R., et al. (2021). A Historical Perspective of Bladderworts (Utricularia): Traps, Carnivory and Body Architecture. Plants 10, 2656. doi:10.3390/plants10122656.

Moseley, H. N. (1884). The fish-eating Utricularia, or bladderwort. Bull. U. S. Fish Comm., 261.

Murali, A., Bhargava, A., and Wright, E. S. (2018). IDTAXA: a novel approach for accurate taxonomic classification of microbiome sequences. Microbiome 6, 140. doi:10.1186/s40168-018-0521-5.

Naseem, F., and Kayang, H. (2018). Fungal endophytes associated with nepenthes khasiana hook. f., An endemic plant of meghalaya, India. Int. J. Curr. Res. Life Sci. 7, 1907–1912.

Okamoto, R., Kojima, H., and Fukui, M. (2017). Acidocella aquatica sp. nov., a novel acidophilic heterotrophic bacterium isolated from a freshwater lake. Int. J. Syst. Evol. Microbiol. 67, 4773–4776. doi:10.1099/ijsem.0.002376.

Oksanen, J., Blanchet, F. G., Friendly, M., Kindt, R., Legendre, P., McGlinn, D., et al. (2022). vegan: Community Ecology Package. Available at: https://cran.r-project.org/package=vegan.

Oliver, D. (1860). Descriptions of New Species of Utricularia from South America, with Notes upon, the Genera Polypompholyx and Akentra. Bot. J. Linn. Soc. 4, 169–176.

Ortega Ardila, A. T., and Romero Salgado, J. O. (2016). Plantas carnívoras de Virolín (Santander, Colombia): una guía de campo.

Peroutka, M., Adlassnig, W., Volgger, M., Lendl, T., Url, W. G., and Lichtscheidl, I. K. (2008). Utricularia: a vegetarian carnivorous plant? Plant Ecol. 199, 153–162. doi:10.1007/s11258-008-9420-3.

Płachno, B. J., Adamec, L., Lichtscheidl, I. K., Peroutka, M., Adlassnig, W., and Vrba, J. (2006). Fluorescence Labelling of Phosphatase Activity in Digestive Glands of Carnivorous Plants. Plant Biol. 8, 813–820. doi:10.1055/s-2006-924177.

Płachno, B. J., Łukaszek, M., Wołowski, K., Adamec, L., and Stolarczyk, P. (2012). Aging of Utricularia traps and variability of microorganisms associated with that microhabitat. Aquat. Bot. 97, 44–48. doi:10.1016/j.aquabot.2011.11.003.

Quast, C., Pruesse, E., Yilmaz, P., Gerken, J., Schweer, T., Yarza, P., et al. (2013). The SILVA ribosomal RNA gene database project: improved data processing and web-based tools. Nucleic Acids Res. 41, D590–D596. doi:10.1093/nar/gks1219.

Quilliam, R. S., and Jones, D. L. (2010). Fungal root endophytes of the carnivorous plant Drosera rotundifolia. Mycorrhiza 20, 341–348. doi:10.1007/S00572-009-0288-4/TABLES/3.

R Core Team (2023). R: A Language and Environment for Statistical Computing. Available at: https://www.r-project.org/.

Richards, J. H. (2001). Bladder function in Utricularia purpurea (Lentibulariaceae): is carnivory important? Am. J. Bot. 88, 170–176. doi:10.2307/2657137.

Rueda-Almazán, J. E., Hernández, V. M., Alcalá-Martínez, J. R., Fernández-Duque, A., Ruiz-Aguilar, M., and Alcalá, R. E. (2021). Spatial and temporal differences in the community structure of endophytic fungi in the carnivorous plant Pinguicula moranensis (Lentibulariaceae). Fungal Ecol. 53, 101087. doi:10.1016/J.FUNECO.2021.101087.

Rutishauser, R. (2016). Evolution of unusual morphologies in Lentibulariaceae (bladderworts and allies) and Podostemaceae (river-weeds): a pictorial report at the interface of developmental biology and morphological diversification. Ann. Bot. 117, 811–832. doi:10.1093/aob/mcv172.

Sanabria-Aranda, L., González-Bermudez, A., Torres, N., Ned Guisande, C., Manjarres-Hernandez, A., Valoyes-Valois, V., et al. (2006). Predation by the tropical plant Utricularia foliosa. Freshw. Biol. 51, 1999–2008. doi:10.1111/j.1365-2427.2006.01638.x.

Sickel, W., Van De Weyer, A. L., Bemm, F., Schultz, J., and Keller, A. (2019). Venus flytrap microbiotas withstand harsh conditions during prey digestion. FEMS Microbiol. Ecol. 95, 10. doi:10.1093/FEMSEC/FIZ010.

Silva, S. R., Gibson, R., Adamec, L., Domínguez, Y., and Miranda, V. F. O. (2018). Molecular phylogeny of bladderworts: A wide approach of Utricularia (Lentibulariaceae) species relationships based on six plastidial and nuclear DNA sequences. Mol. Phylogenet. Evol. 118, 244–264. doi:10.1016/j.ympev.2017.10.010.

Singh, K., Reyes, R. C., Campa, G., Brown, M. D., Hidalgo, F., Berg, O., et al. (2020). Suction Flows Generated by the Carnivorous Bladderwort Utricularia—Comparing Experiments with Mechanical and Mathematical Models. Fluids 5, 33. doi:10.3390/fluids5010033.

Sirová, D., Adamec, L., and Vrba, J. (2003). Enzymatic activities in traps of four aquatic species of the carnivorous genus Utricularia. New Phytol. 159, 669–675. doi:10.1046/j.1469-8137.2003.00834.x.

Sirová, D., Bárta, J., Borovec, J., and Vrba, J. (2018a). The Utricularia-associated microbiome: composition, function, and ecology. Oxford University Press doi:10.1093/oso/9780198779841.003.0025.

Sirová, D., Bárta, J., Šimek, K., Posch, T., Pech, J., Stone, J., et al. (2018b). Hunters or farmers? Microbiome characteristics help elucidate the diet composition in an aquatic carnivorous plant. Microbiome 6, 225. doi:10.1186/s40168-018-0600-7.

Sirová, D., Borovec, J., Černá, B., Rejmánková, E., Adamec, L., and Vrba, J. (2009). Microbial community development in the traps of aquatic Utricularia species. Aquat. Bot. 90, 129–136. doi:10.1016/j.aquabot.2008.07.007.

Taylor, P. (1989). The genus Utricularia. A taxonomic monograph. Kew Bull., 1–724.

Valdés, C. M. P. (2008). El género Utricularia (lentibulariaceae) en las antillas mayores. Rev. del Jardín Botánico Nac. 29, 11–20.

VanInsberghe, D., Maas, K. R., Cardenas, E., Strachan, C. R., Hallam, S. J., and Mohn, W. W. (2015). Non-symbiotic Bradyrhizobium ecotypes dominate North American forest soils. ISME J. 9, 2435–2441. doi:10.1038/ismej.2015.54.

Walker, I. (2004). Trophic interactions within the Utricularia habitat in the reservoir of the Balbina hydroelectric powerplant(Amazonas, Brazil). Acta Limnol. Bras. 16, 183–191.

Wang, Q., Garrity, G. M., Tiedje, J. M., and Cole, J. R. (2007). Naive Bayesian classifier for rapid assignment of rRNA sequences into the new bacterial taxonomy. Appl Env. Microbiol 73, 5261–5267. doi:10.1128/AEM.00062-07.

Weiss, J. V., Rentz, J. A., Plaia, T., Neubauer, S. C., Merrill-Floyd, M., Lilburn, T., et al. (2007). Characterization of Neutrophilic Fe(II)-Oxidizing Bacteria Isolated from the Rhizosphere of Wetland Plants and Description of Ferritrophicum radicicola gen. nov. sp. nov., and Sideroxydans paludicola sp. nov. Geomicrobiol. J. 24, 559–570. doi:10.1080/01490450701670152.

Westermeier, A. S., Fleischmann, A., Müller, K., Schäferhoff, B., Rubach, C., Speck, T., et al. (2017). Trap diversity and character evolution in carnivorous bladderworts (Utricularia, Lentibulariaceae). Sci. Rep. 7, 12052. doi:10.1038/s41598-017-12324-4.

Wu, L., Han, T., Li, W., Jia, M., Xue, L., Rahman, K., et al. (2013). Geographic and tissue influences on endophytic fungal communities of taxus chinensis var. mairei in China. Curr. Microbiol. 66, 40–48. doi:10.1007/S00284-012-0235-Z/FIGURES/2.

Zahran, H. H. (1999). Rhizobium -Legume Symbiosis and Nitrogen Fixation under Severe Conditions and in an Arid Climate. Microbiol. Mol. Biol. Rev. 63, 968–989. doi:10.1128/MMBR.63.4.968-989.1999.

